# Virome analysis provides new insights into the pathogenesis mechanism and treatment of SLE disease

**DOI:** 10.1101/2023.10.25.564084

**Authors:** Yifan Wu, Zhiyuan Zhang, Xun Liu, Ye Qiu, Xingyi Ge, Zhichao Miao, Xiangxian Meng, Yousong Peng

## Abstract

Systemic lupus erythematosus (SLE) is a multi-system autoimmune disease. Many viruses have been reported to be associated with SLE, but the diversity of the virome in SLE and the mechanisms underlying the interactions between viral infection and SLE are still unclear. This study identified ten human virus species from 826 RNA-Seq samples of human blood from 688 SLE patients and 138 healthy controls. SLE patients were found to have higher positive rates of viruses than healthy controls, although the virus abundances were low and comparable in both SLE patients and healthy controls. Analysis of the antiviral interferon-stimulated genes (ISGs) in samples showed higher ISG expression levels in virus-positive samples compared to virus-negative samples, which confirmed viral infections in virus-positive samples. Further analysis of differential expression genes between virus-positive and negative SLE patients showed that several genes that were up-regulated in SLE patients were further up-regulated after viral infections, and they were mainly enriched in immune response-related biological processes, suggesting an excess immune response in SLE patients after viral infections. Finally, three marker genes of the SLE severity, namely STAT1, STAT5A, and STAT5B, showed higher expression levels in virus-positive SLE patients compared to those in virus-negative SLE patients, suggesting that viral infections may aggravate the SLE disease. Overall, the study deepens our understanding of the association between viruses and SLE and provides new insights into prevention and control of the disease.

## Introduction

Systemic lupus erythematosus (SLE) is a multi-system autoimmune disease with most patients being women of childbearing age. The incidence rate in women is approximately ten times higher than that in men^1^. SLE is characterized by the simultaneous dysregulation of innate and adaptive immune systems, leading to the production of pathogenic autoantibodies and activation of the type I interferon pathway^2,3^. Additionally, hematological complications, such as anemia, leukopenia, and thrombocytopenia, are frequently observed in SLE patients. These symptoms may result from either bone marrow failure or excessive peripheral cell destruction, both of which are immune-mediated^4^. Drugs and infections are also common causes of SLE. Although several genetic regions have been reported to be associated with SLE, the pathogenesis mechanism of the SLE disease remains unclear^5^.

Viruses have been shown to modify the clinical picture of several autoimmune diseases, such as SLE, type 1 diabetes, rheumatoid arthritis, Sjögren’s syndrome, herpetic stromal keratitis, celiac disease, and multiple sclerosis^6-8^. Infections of multiple viruses including Coxsackie B virus (CVB), rotavirus, influenza A virus (IAV), and herpesvirus, have been proposed to modulate the induction and development of autoimmune diseases^9,10^. Many studies have investigated the relationship between SLE and viruses. For example, Stearrett et al. identified differentially expressed human endogenous retroviruses in SLE patients^11^; Pan et al. reported strong associations between SLE and several viruses including Epstein-Barr virus (EBV or HHV4), parvovirus B19 (B19V) and cytomegalovirus (CMV or HHV5)^12^; Guo et al. identified a large number of viruses in the peripheral blood mononuclear cells (PBMCs) of 10 SLE patients^13^. Despite these studies, the virome diversity in the SLE disease and the mechanisms underlying the interactions between viral infection and SLE are still unclear.

With the development of next-generation-sequencing technology, virome studies have identified many viruses in humans. For example, Kumata et al. identified 39 animal viruses in 51 somatic tissues of healthy people by analyzing 8991 samples from the GTEx project^14^. Ye et al. built the Human Virus Database which included more than 1000 animal viruses observed in 68 human tissues^15^. Although virome diversity has been extensively explored, viral contributions to human health and disease are still often overlooked. For example, lots of evidence indicated that persistent human cytomegalovirus infection is associated with atherosclerosis and coronary artery disease^16^; the severe acute respiratory syndrome coronavirus 2 (SARS-CoV-2) has been reported to potentially cause multiple diseases such as placenta diseases, endocarditis, memory disorders and so on^17-20^. Thus, it is crucial to identify viruses associated with human diseases to better understand the pathogenic mechanisms underlying these diseases. For example, Kim et al. identified two viruses (picobirnavirus and tobamovirus) which were more prevalent in pregnant women with T1D than in healthy ones^21^; Deng et al. identified 9 viruses in brain tissues of the Parkinson’s disease (PD) patients and found higher positive rate of most viruses in PD patients compared to healthy individuals^22^. Unfortunately, most published virome studies on human diseases do not include further investigations into the interactions between the virome and the diseases.

This study aimed to investigate the virome diversity of the SLE disease and the association between viral infections and the disease. First, we collected a large number of SLE-related blood transcriptome datasets from public databases and identified the virome from the data; second, the virome was validated by the analysis of the expression level of antiviral interferon-stimulated genes (ISGs) in samples; third, we identified human genes related to the interaction between SLE and viral infections and analyzed their functions; finally, we analyzed the expression levels of several marker genes related to the severity of SLE in SLE samples. Overall, this study systematically investigated the virome in the SLE disease and the complex interaction between viral infection and the disease, which provides new insights into the pathogenesis mechanisms of SLE.

## Materials and Methods

### Data Collection

The SLE-related RNA-Seq data were retrieved from the NCBI Sequence Read Archive (SRA) database by the steps described as follows. Firstly, we searched the SRA database using “SLE” or “systemic lupus erythematosus” as terms and kept all transcriptome sequencing samples related to SLE, which resulted in 44 projects and 2,020 runs. Subsequently, we conducted a manual inspection of all samples, retaining only those derived from blood tissue, resulting in 826 samples from 15 projects, which included 688 and 138 samples from SLE patients and healthy controls, respectively (Table S1).

### Virus identification

A computational workflow was developed to identify viruses from the RNA-Seq data. Firstly, fastp (version=0.20.0) was used to trim adapters and filter low-quality reads. The remaining reads were aligned to the human genome hg38 with bwa (version=0.7.12-r1039). We then queried the unaligned reads using BLASTN (version=2.5.0) against a library of genomic sequences from animal viruses with hosts registered as invertebrates, vertebrates, or humans. The reads with an E-value of less than 1E-10 for the best hit were labeled as hypothetical viral reads. Finally, the putative viral reads were queried against the NT database (downloaded on October 12^th^, 2022) with BLASTN to remove false positives. We retained reads with the best match to animal viruses as true viral reads.

### Filtering low-reliability viral reads

The low-reliability viral reads were further filtered in the following three aspects: Firstly, the viruses with fewer than three reads mapped and detected only in one dataset were removed; secondly, we removed viruses that have not been reported to infect humans; thirdly, viruses of the Retroviridae family were removed because there is a possibility of cross-contamination from endogenous retroviral sequences, and viruses of the Baculoviridae family were also removed as they are commonly used in the laboratory^15^.

### Virus depth calculation

Since the RNA-Seq data used in this study were not originally designed for virus discovery, which could result in an underestimation of virus abundance in humans, the viral abundance in a sample was normalized as 1000 times of Reads Per Kilobase per Million mapped reads (RPKM) according to our previous study^22^. Specifically, it was calculated as follows:

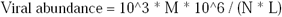

where M is the number of RNA-Seq reads assigned to a certain virus, N is the total number of RNA-Seq reads assigned to the human genome, and L is the length of the viral genome in base pairs.

### Differential expression analysis of human genes

Raw sequencing reads were quality-controlled as previously described. The resulting clean data were mapped to the human genome hg38 using bwa (version=0.7.12-r1039) with default parameters, and BAM files were sorted with Samtools (version=1.9). Gene counts were obtained using the featureCounts program (version= 2.0.1). The batch-corrected gene expression matrix was generated using the “removeBatchEffect” function in the limma package (version=3.46.0) in R. Differentially expressed genes (DEGs) between two groups were identified using the “DESeq” function of the DESeq2 package (version=1.30.1) in R. Genes with at least two-fold changes of expressions and the adjusted p-value < 0.05 were considered as DEGs.

### Functional enrichment analysis

Functional enrichment of human genes was performed using the Gene Ontology (GO) and Kyoto Encyclopedia of Genes and Genomes (KEGG) pathway analysis by the clusterProfiler package (version=3.18.1) in R. All KEGG pathways and GO terms with FDR adjusted p-values less than 0.05 were considered significant enrichment.

### Statistical analysis

All statistical analyses were conducted in Python (version 3.7.9). The Wilcoxon Rank Sum Test was conducted using the “scipy.stats.ranksums” function.

## Results

### Overall workflow of the study

The workflow of the study is shown in Figure 1 and contains three sections. The first section was “Dataset Retrieval”, during which 826 samples of human blood including 688 from SLE patients and 138 from healthy controls with RNA-Seq data were obtained from 15 datasets in the NCBI SRA database. The second section was “Virus Identification”, during which human viruses were identified from these samples using a homology-based method, and their abundances were quantified based on the RNA-Seq data. The third section, titled “Host Gene Analysis”, involved quantifying human gene expression profiles and using them to analyze ISG expressions, the interactions between SLE and viral infections, and SLE severity.

**Figure 1.**
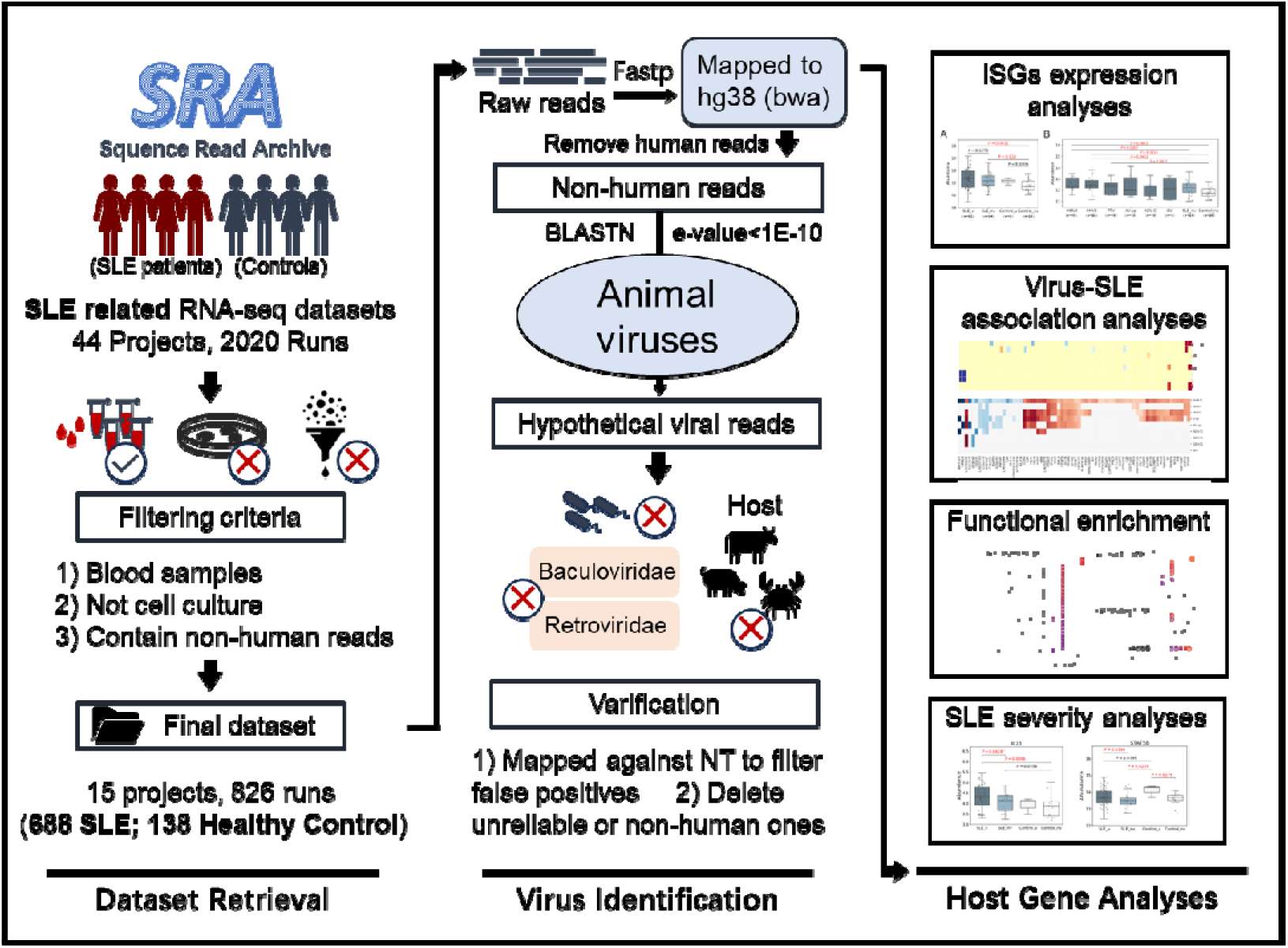
Graphical workflow of the study. It contained three sections: Dataset Retrieval, Virus Identification and Host Gene Analysis.

### Ten human viruses were identified with low abundances and low positive rates

Among all 826 human blood samples mentioned above, a total of 10 human viruses, including three viruses from the Herpesviridae family (Human alphaherpesvirus 3 (HHV3), Human gammaherpesvirus 4 (HHV4), Human betaherpesvirus 5 (HHV5)), two viruses from the Anelloviridae family (Torque teno virus (TTV), Anelloviridae sp. (AV-sp)), two viruses from the Flaviviridae family (Human pegivirus (PGV-A), GB virus C (PGV-C)), one virus from the Adenoviridae family (Human mastadenovirus C (HAdV-C)), one virus from the Orthomyxoviridae family (Influenza A virus (IAV)) and one virus from the Picornaviridae family (Cardiovirus A (EMCV)), were identified in 84 SLE patients and 7 healthy controls (Figure 2A & Table S2). Most virus-positive samples (n=76) contained only one virus; in the remaining ones (n=8), 2∼4 viruses were identified. Half of the viruses were identified in fewer than 10 samples. HHV5 was the most commonly detected virus and was identified in 28 samples, accounting for 3.4% of all samples, while EMCV was only detected in one sample (Figure 2B). Analysis of the virus abundance in blood samples showed that the majority of viruses had low abundances, with the exception of pegivirus (PGV-A and PGV-C in Figure 2C), which has been reported to be widely distributed in human populations and has long-term infections in humans^23^.

**Figure 2.**
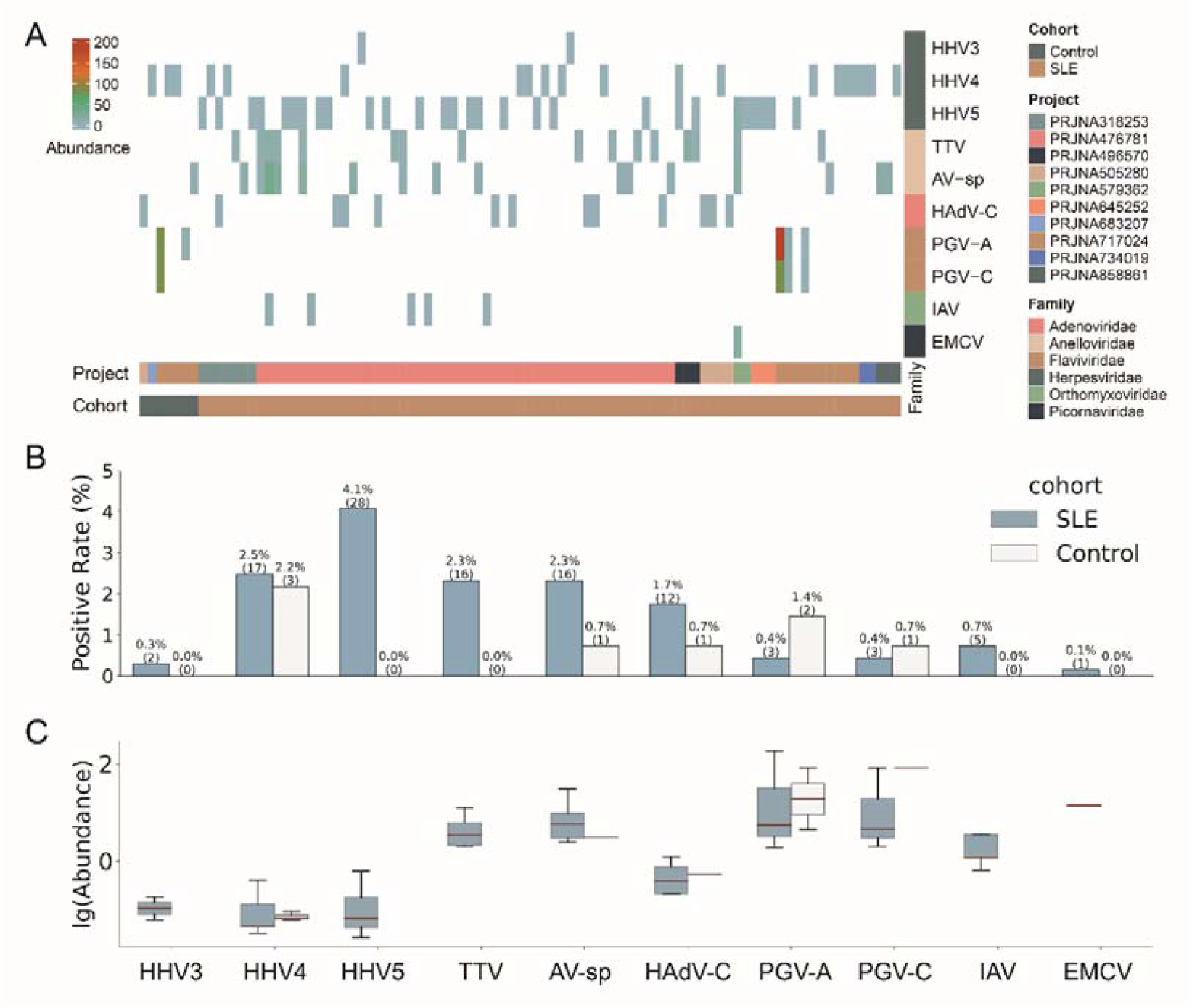
Overview of the virome identified in the study. (A) The abundance of viruses identified in positive samples. Each column represents a sample with the project and cohort information tagged by the color bars at the bottom. (B) The comparison of virus positive rates between SLE and Control groups. The numbers of positive samples were tagged in the brackets. (C) The comparison of virus abundance between SLE and Control groups. For clarity, virus names were shown as abbreviations. The full names and taxonomy IDs of these viruses were shown in Table S3.

### SLE patients had higher positive rates of viruses than healthy controls

We then compared the viromes identified from SLE patients and healthy controls. All ten viruses mentioned above were identified in SLE patients, while only five of them (HHV4, AV-sp, HAdV-C, PGV-A, and PGV-C) were identified in healthy controls. Regarding positive rates, all viruses, except for PGV-A and PGV-C, had higher rates in SLE patients than in healthy controls, although the positive rates of all viruses were less than 5% (Figure 2B). For example, both HHV5 and TTV were only detected in SLE patients with a positive rate of 4.1% and 2.3%, respectively. When taking all viruses together, SLE patients exhibited a virus-positive rate more than twice that in healthy controls (12.2% vs 5.1%). Most viruses had similar or lower abundances in SLE patients than in healthy controls. Interestingly, both PGV-A and PGV-C had higher positive rates and abundances in healthy controls than in SLE patients (Figure 2B and 2C). To sum up, viruses identified from SLE patients showed greater diversity and higher positive rates than those from healthy controls.

### Higher ISG expression levels in virus-positive samples

The Type I interferon (IFN-I) response is crucial in combating viral infections, primarily by inducing the expression of ISGs that possess antiviral functions^24^. It also plays an important role in the development of SLE disease, and the expression of certain ISGs is increased in SLE patients^25^. To validate the virome identified in the study, a set of antiviral ISGs were obtained from Zhou’s study^26^, and a set of SLE-related ISGs were obtained from Siddiqi’s study (Table S4)^27^. As expected, the expression level of SLE-related ISGs were higher in SLE patients than in non-SLE patients (Figure S1). Then, the SLE-related ISGs were excluded from the antiviral ISGs to remove the influence of SLE on ISG expressions, leading to a set of non-SLE-related antiviral ISGs. The expression levels of non-SLE-related antiviral ISGs were analyzed to confirm the presence of viral infection in virus-positive samples. As shown in Figure 3A, the non-SLE-related antiviral ISGs had slightly higher expressions in virus-positive samples than in virus-negative ones. Further analysis of these genes in virus-positive samples grouped by virus (Figure 3B) showed that for all viruses except AV-sp, the median expression levels of non-SLE-related antiviral ISGs in virus-positive groups were higher than those in virus-negative Control group, although statistical differences were only observed for HHV4 (p-value=0.0453) and HHV5 (p-value=0.0377).

**Figure 3.**
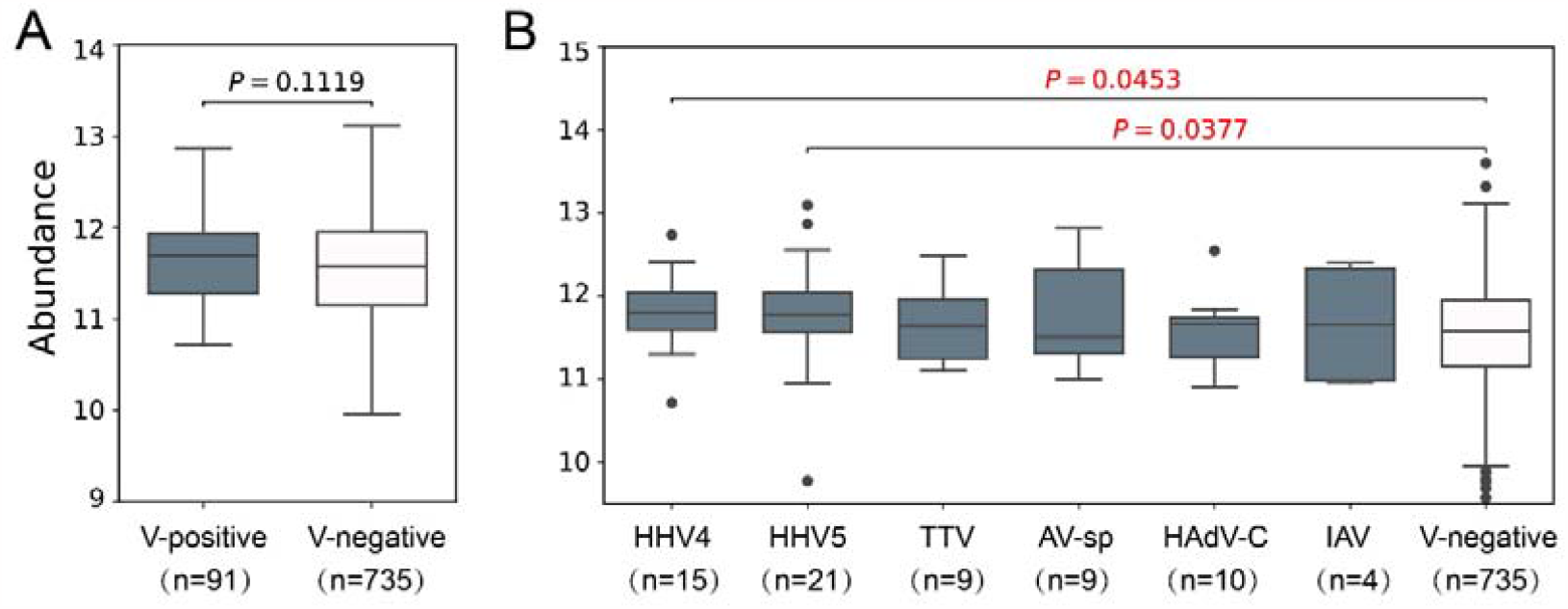
Analysis of expression level of non-SLE-related antiviral ISGs in samples used in the study. (A) The comparison of expression levels of non-SLE-related antiviral ISGs between virus-positive samples and virus-negative samples. (B) The comparison of expression levels of non-SLE-related antiviral ISGs in virus-positive samples grouped by virus to that in virus-negative samples. *P-*values based on Wilcoxon rank sum test were tagged. Samples which were found to contain two or more viruses were excluded in the analysis. Only six viruses were analyzed as less than 3 samples were positive for the other four viruses. The numbers of samples in each group were shown in brackets.

### The immune system was over-activated in SLE patients after viral infections

Then, the interactions between the SLE disease and viral infections were investigated. Firstly, the differentially expressed human genes in virus-negative SLE samples compared to virus-negative control samples were identified to investigate the influence of the SLE on human gene expressions. A total of 463 up-regulated and 52 down-regulated DEGs were identified and defined as SLE-related genes (Figure 4A). The up-regulated SLE-related genes were mainly enriched in terms related to humoral immunity, immunoglobulin and phagocytosis, with “humoral immune response” being the most enriched biological process (Figure 4B). Meanwhile, no significantly enriched functions were obtained for the down-regulated SLE-related genes.

**Figure 4.**
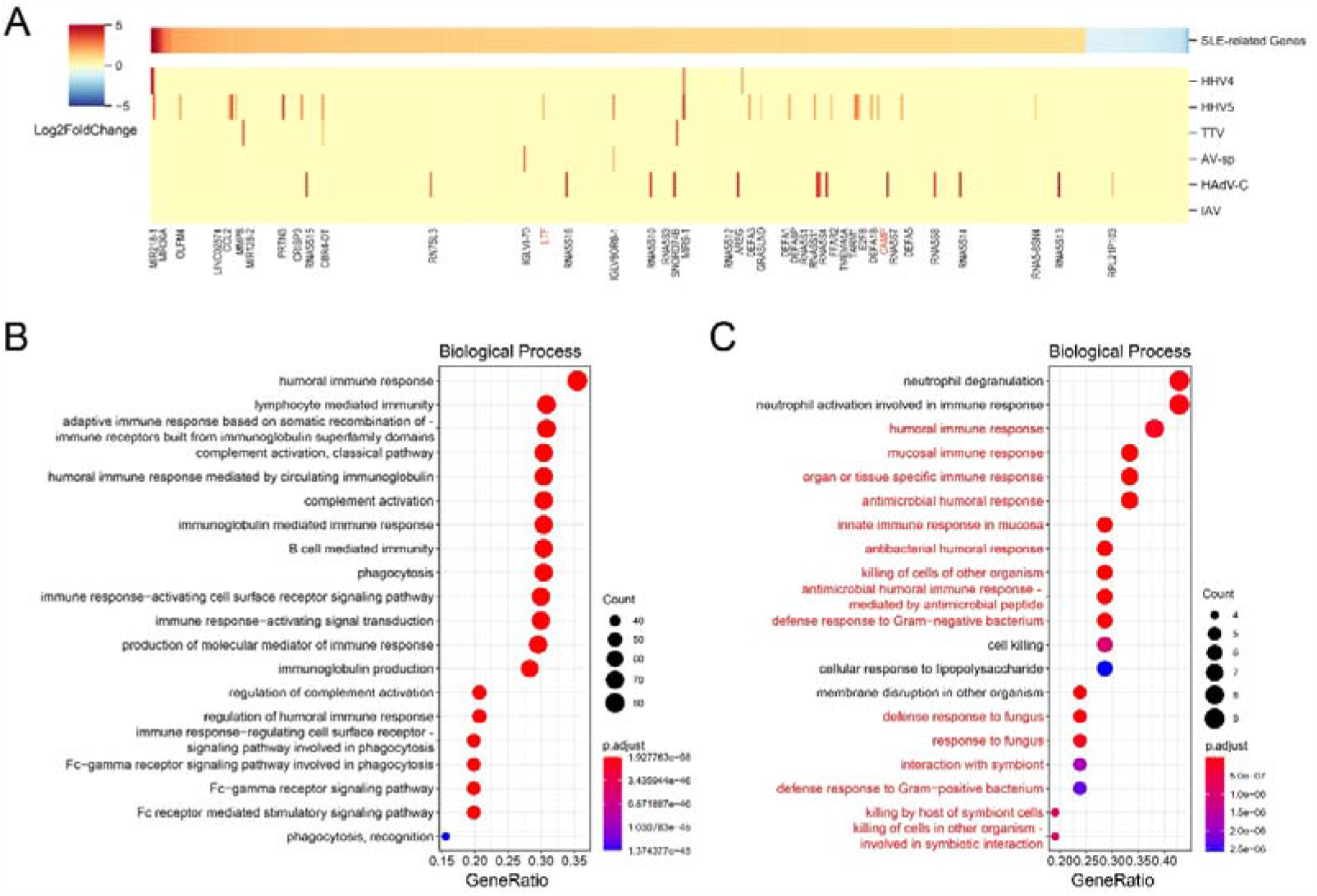
Analyses of the interactions between the SLE disease and viral infections. (A) Differential analysis of SLE-related genes between virus-positive SLE samples and virus-negative SLE samples. Samples which were found to contain two or more viruses were excluded in the analysis. Only six viruses were analyzed as less than 3 virus-positive samples were available for other four viruses. (B) Top 20 enriched biological processes for up-regulated SLE-related genes. (C) Top 20 enriched biological processes for further up-regulated SLE-related genes in virus-positive SLE samples. The biological processes that were related to immune responses and were also enriched in all SLE-related genes were highlighted in red.

To investigate the influence of viral infections on the SLE disease, for each virus, the DEGs in virus-positive SLE samples compared to virus-negative SLE samples were identified and were defined as Virus-SLE-Interaction-Genes (VSIGs). Four viruses, including HHV3, EMCV, PGV-A and PGV-C, were not included in the analysis as less than 3 virus-positive samples were available for them. As shown in Figure 4A, expression levels of most SLE-related genes were not significantly changed by viral infections. Interestingly, for the 42 genes with significant changes, all of them were up-regulated after viral infections. For example, two immune-related genes, CAMP and LTF (highlighted in red in Figure 4A), both of which were up-regulated in SLE patients, were further up-regulated after viral infections. Analysis of functions of these genes showed that they were mainly enriched in biological processes related to immune responses, such as “neutrophil degranulation”, “neutrophil activation involved in immune response”, “humoral immune response”, and so on. 3/4 of these enriched biological processes were also observed for the SLE-related genes (highlighted in red in Figure 4C).

The number of VSIGs varied much among viruses and ranged from 31 to 900. The HHV5 had the largest number of VSIGs, including 556 up-regulated and 344 down-regulated VSIGs, while the HHV4 had the fewest VSIGs, including only 13 up-regulated and 18 down-regulated VSIGs. Interestingly, most of VSIGs of HAdV-C were up-regulated, while most of VSIGs of TTV were down-regulated. Analysis of the overlapped VSIGs between viruses showed that few VSIGs were shared between viruses (Figure S2).

### Viral infection aggravated the SLE

It is suspected that the SLE patients may be aggravated after viral infections since the immune system was over-activated after viral infections. Therefore, two SLE severity marker genes, i.e., STAT1 and STAT5, elevated expressions of which in CD4+ T cells or B cells were associated with aggravation of SLE disease, were collected from studies of Goropevšek et al. and Aue et al. ^28,29^. As expected, the expression level of STAT1, STAT5A and STAT5B (both of latter two genes also known as STAT5) were higher in the virus-positive SLE group than those in the virus-negative SLE group (Figure 5A), with significant differences observed for STAT1 (p-value = 0.0125) and STAT5B (p-value = 0.0059). Analysis of the expression levels of these marker genes by virus showed that although the expression level of different marker genes varied among virus-positive samples of different viruses, in most of the time, virus-positive SLE samples had higher expression levels of marker genes compared to the virus-negative SLE group (Figure 5B).

**Figure 5.**
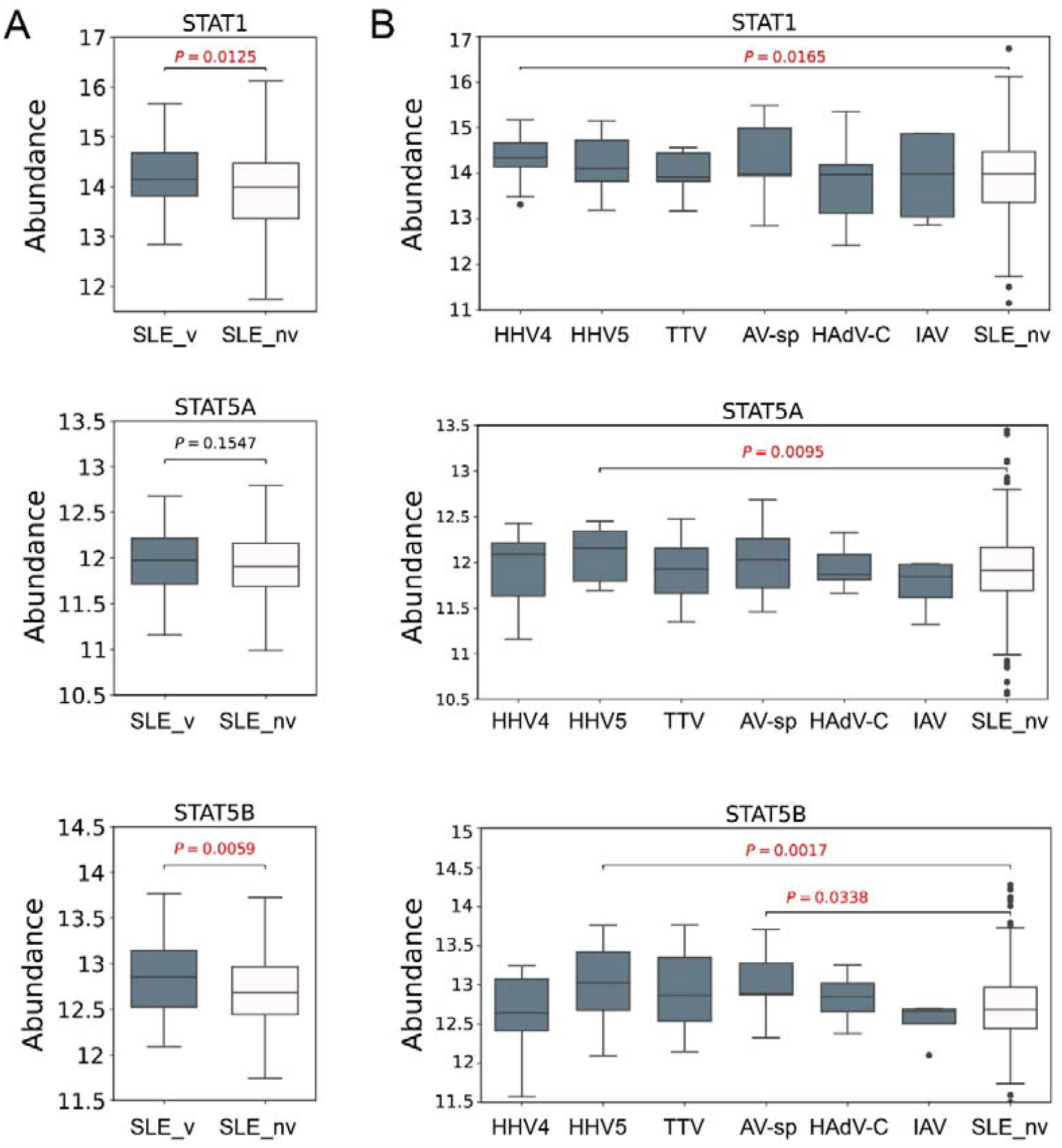
The expression level of SLE severity marker genes. (A) The expression levels of STAT1, STAT5A and STAT5B, respectively, in virus-positive (SLE_v) and virus-negative SLE patients (SLE_nv). (B) The expression levels of STAT1, STAT5A and STAT5B, respectively, in specific-virus-positive SLE patients and virus-negative SLE patients. Figures were titled by the gene names. *P-*values based on Wilcoxon rank sum test were tagged.

## Discussion

In this study, we collected the most abundant blood transcriptome sequencing data of the SLE disease so far, and conducted rigorous identification of human viruses, obtaining a comprehensive virus profile in the blood of SLE patients. A total of ten human virus species were detected with low positive rates and low abundances in SLE patients; however, the positive rates of viruses were higher in the SLE patients than those in healthy controls, suggesting that the SLE patients may be more susceptible to viral infections than healthy people, which was consistent with previous studies^30^. Analysis of antiviral ISG expression level confirms viral infections in virus-positive samples, further validating the virome identified in our study.

Previous studies by Guo et al. identified 36 human-associated viruses (not at the species level) from the RNA-Seq data of the PBMCs of 10 SLE patients^13^. In contrast, our study only identified ten human virus species from more than 600 samples of SLE patients. The difference was mainly due to the different workflows used for virus identification. Guo et al. identified viruses by firstly removing all reads aligned to the human genome and then by querying against RefSeq virus genomes with the unmapped reads using BLASTN. All reads that had an E-value smaller than 1E-5 in the BLAST were taken as viral reads. Compared to their workflow, we used a more reliable strategy (Materials and Methods): i) only reads which had an E-value of smaller than 1E-10 in the BLAST were taken as hypothetical viral reads; ii) the hypothetical viral reads were further queried against the NCBI NT database to remove false positives; iii) low-reliability viruses were further removed due to low abundance or potential contamination. Besides, viral infections of samples were confirmed by analysis of the ISG expressions. Thus, although far fewer viruses were identified, the virome identified in our study was much more reliable than that in Guo’s study.

The ten virus species identified in our study have been commonly reported in humans: i) Three viruses from the Herpesviridae family (HHV3, HHV4, and HHV5) have frequently been reported in humans with a high positive rate^9,31,32^. Among them, HHV4 is considered to be definitely associated with SLE^12^, and researchers have observed elevated SLEDAI (Systemic Lupus Erythematosus Disease Activity Index) scores in patients with lytic HHV4 infections^33^. HHV5 is also believed to contribute significantly to the development of SLE, as multiple studies have indicated a higher prevalence of HHV5 among SLE patients^12,34^. Notably, our identification results revealed that HHV4 and HHV5 exhibited the highest positive rate in SLE patients, with HHV5 exclusively detected in SLE samples. This further substantiates their strong association with SLE. Subsequent analyses focusing on ISG expression and markers of SLE severity highlighted the most prominent differences between these two virus-positive samples and control samples, thus reinforcing the substantial impact of these viral infections on SLE. ii) TTV and AV-sp, which exhibited relatively low positive rates, are categorized as non-pathogenic anelloviruses that are believed to be widely present in the human blood virome^23^. It is noteworthy that multiple studies have reported a higher positive rate of anelloviruses in SLE patients compared to that in healthy people^30,35^. Our results concerning VSIGs suggested that TTV, compared to other viruses, had a highly similar expression profile with HHV5 after viral infection of SLE patients (Figure S2), implying a possible similar mechanism of their influences on SLE. Interestingly, the expression of one marker gene (STAT5B) associated with SLE severity was also significantly higher in AV-sp positive samples compared to the virus-negative SLE patients, suggesting its possible associations with SLE, which necessitates further investigation. iii) Two members of the Flaviviridae family, PGV-A and PGV-C, are also generally regarded as non-pathogenic. Compared to anelloviruses, most researchers believe that their population distribution is relatively smaller^23^, which is consistent with our results. iv) HAdV-C was indicated by a higher prevalence in SLE patients in our results, despite its low detection rate in the general population^36^. v) As for IAV and EMCV, limited reports suggest their potential connection with autoimmune diseases, although the IAV is one of the most prevalent respiratory viruses in human populations^37^.

SLE patients have concurrent dysregulation of innate and adaptive immune systems^2^. Our study revealed the increased expressions of SLE-related ISGs in SLE patients. Most SLE-related genes were up-regulated and primarily enriched in immune-related biological processes. Interestingly, certain up-regulated genes in SLE patients were further up-regulated after viral infections, primarily involved in immune responses. This finding aligns with a previous study by Han et al., which reported the evaluated expression of specific ISGs in SLE patients following HHV4 infections^33^. This suggests that viral infections may trigger an over-activated immune response, potentially exacerbating the SLE disease. This was partially validated by our findings, which showed that three severity marker genes of SLE had higher expressions in virus-positive SLE samples than in virus-negative SLE samples. These findings suggest the importance of timely monitoring the viral infection status of SLE patients and preventing viral infections during the treatment of the disease. This proactive approach may effectively reduce the likelihood of rapid disease progression or deterioration.

There were some limitations to the study. Firstly, only 688 SLE samples were used in the analysis, and the RNA-Seq data of these samples in the original studies were not designed for virus discovery, which may lead to the underestimation of the virus diversity and abundance in SLE diseases. Many more studies are needed to uncover the virome diversity in SLE disease. Secondly, when analyzing differential gene expression, it is difficult to separate the effect of SLE disease and the infection of specific viruses, as there were few virus-positive healthy controls for each virus. Further investigations with more blood samples may improve the study.

## Conclusions

This is the first study to systematically investigate the virome in the SLE disease and the complex interaction between viral infections and the disease. Our findings contribute to the understanding of the association mechanism between SLE disease and viral infections, and also provide new clues for the prevention and control of the disease.

## Acknowledgements

We thank members in PengLab for helpful discussions on the manuscript.

## Availability of data and materials

All data used in the study was provided in supplementary materials.

