## Supplementary Figures for "Virome analysis provides new insights into the pathogenesis mechanism and treatment of SLE disease"

**Figure S1.** The comparison of SLE-related ISGs between SLE samples and non-SLE samples. *P-*values based on Wilcoxon rank sum test were tagged. The numbers of samples in each group were shown in brackets.

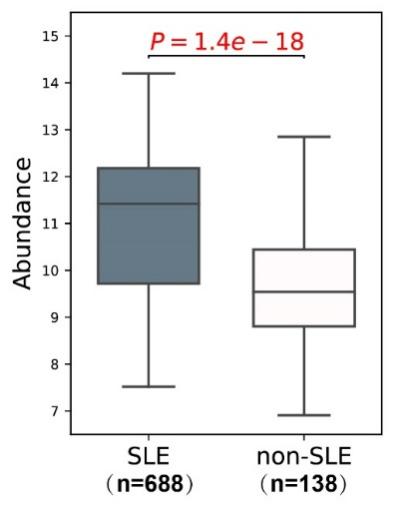

**Figure S2.** The numbers of overlapped up-regulated (A) and down-regulated (B) VSIGs between viruses. Samples which were found to contain two or more viruses were excluded from the analysis. Only six viruses were analyzed as less than 3 virus-positive samples were available for the other four viruses.

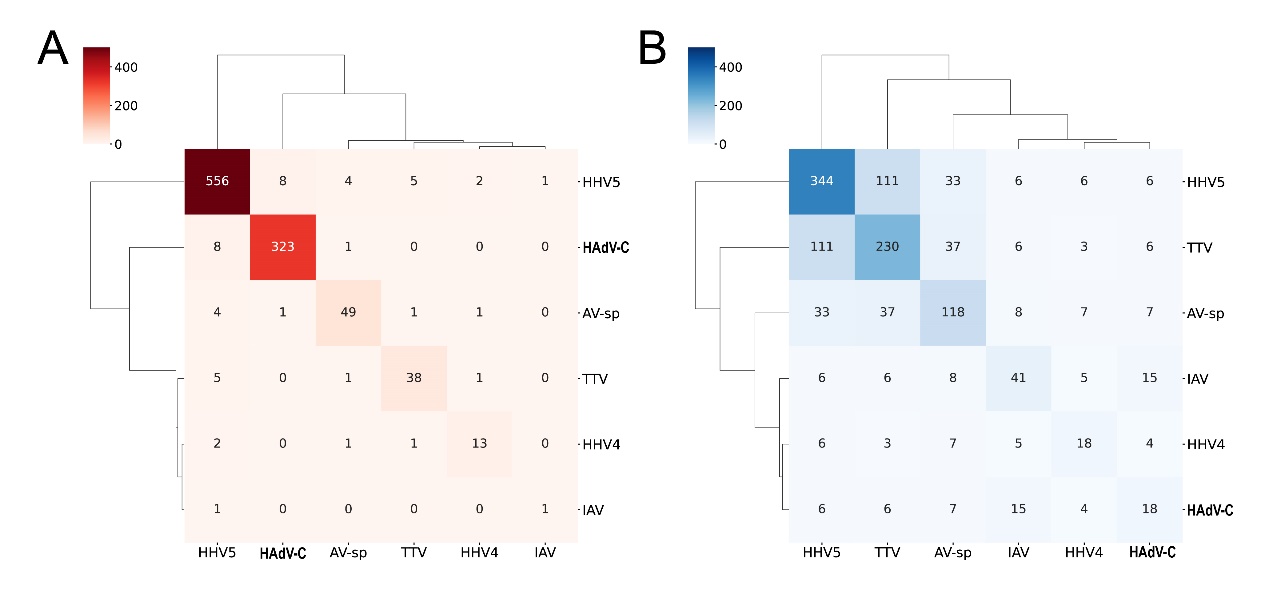
